# Wine yeast phenomics: a standardized fermentation method for assessing quantitative traits of *Saccharomyces cerevisiae* strains in enological conditions

**DOI:** 10.1101/191353

**Authors:** Emilien Peltier, Margaux Bernard, Marine Trujillo, Duyên Prodhomme, Jean-Christophe Barbe, Yves Gibon, Philippe Marullo

**Affiliations:** Univ. Bordeaux, ISVV, Unité de recherche OEnologie EA 4577, USC 1366 INRA, Bordeaux INP, Villenave d’Ornon, France.; Biolaffort, Bordeaux, France.; Pernod Ricard, Creteil, France; INRA, University of Bordeaux, UMR 1332 Fruit Biology and Pathology, F-33883 Villenave d’Ornon, France

## Abstract

This work describes the set up of a small scale fermentation methodology for measuring quantitative traits of hundreds of samples in an enological context. By using standardized screw cap vessels, the alcoholic fermentation kinetics of *Saccharomyces cerevisiae* strains were measured by following the weight loss over time. Preliminary results showed that the kinetic parameters measured are in agreement with those observed in larger scale vats. The small volume used did not impair any analysis of the strain performance. Indeed, this fermentation system was coupled with robotized enzymatic assays and 8 end-point metabolites of enological interest were measured accurately. Moreover, the vessel used offers the possibility to assay 32 volatiles compounds using a headspace solid-phase microextraction coupled to gas chromatography-mass spectrometry approach. Data presented demonstrates that the shaking conditions significantly impacted the mean and the variance of kinetic parameters, primary metabolites, and the production of volatile compounds. This effect was likely due to an enhanced transfer of dissolved oxygen during the first hours of the alcoholic fermentation. To test the efficiency of this experimental design, the phenotypic response of 35 wine industrial starters was measured in five grape juices from the Bordeaux area. A multivariate analysis illustrated that strains were better discriminated by some grape must, than others. The technological performances as well as the phenotypic robustness of these strains was measured and discussed. The optimized methodology developed allows investigating multiple fermentation traits for numerous yeast strains and conditions and greatly contributes in achieving quantitative genetics studies as well as yeast selection programs.

## Introduction

In the last decade, the emergence of NGS (Next Generation Sequencing) has opened perspectives for studying the genetic adaptation of microbial species in their environments [1]. This is the case for the wine microbiome [2,3], which is subjected to a complex and evolving environment from grape must to wine. Thanks to the reduction in the genome sequencing costs, large comparative genomic studies were carried out at the intraspecific level for lactic bacteria (*Oenococcus oeni)* [4] and various yeast species, including *S. uvarum* [2,3], *B. bruxellensis* [5,6] and *S. cerevisiae* [7]. The bioinformatics analysis of such genomes shed light on genomic adaptation mechanisms such as chromosomal introgression [5], chromosomal translocations [8,9], horizontal transfer [10,11], polyploidy [5,6]; for an extensive review see [12]. Moreover, since most of those species are found in other environments, population structure studies based on SNP analysis clearly demonstrated the wine microbial domestication in link with its environmental origin [4,5,13,14].

To bridge the gap existing between all this diversity and the molecular mechanisms of phenotypic adaptation, functional genetics studies have to be achieved. In order to decipher the molecular basis of phenotypic diversity, quantitative genetics approaches such Quantitative Trait Loci (QTL) mapping or Genome Wide Association Studies (GWAS) can be used [15]. QTL mapping turns out to be particularly efficient for identifying natural genetic variations controlling relevant traits in enology [8,16–21].

One of the main limitations of this approach is the requirement of intensive genotyping and phenotyping work. While the genotyping task can be easily achieved with NGS strategies [22,23], the measurement of complex phenotypes for several hundreds of individuals is not yet an easy task. Recently, various methods for measuring yeast phenotypes in a high troughtput way has been reviewed and reffered to as *phenomics* [24]. Although very efficient and standardized, these methods are mostly used for measuring yeast fitness (growth) but partially fail to measure the fermentation performance. Indeed, physiological studies showed that during the stationary growth phase, huge phenotypic discrepancies can be measured among strains having similar growth parameters [25]. Moreover, individuals showing the best growth are not always the most efficient during the fermentation [11,26]. Beyond the fermentation rate, the measurement of other phenotypes is critical. In fact, during fermentation, yeasts produce and/or consumes compounds, that affect the organoleptic qualities of the resulting wine [17,27,28]. Therefore, standardized methods for measuring wine fermentation phenotypes are required.

In this work, we set up a methodology for phenotyping several enological traits in 10 mL-vials with a good reliability. The effect of shaking was particularly investigated and strongly impacted the phenotypic response of yeast. The phenotypic characterization of 35 industrial starters was measured in 5 different grape musts, highlighting some interesting genetics *x* environmental interactions.

## Materials and Methods

### Yeast strains and culture media used

All the yeast strains used belong to the *Saccharomyces cerevisiae* species. Four strains are monosporic clones derived from industrial wine starters that have been previously described [18,20]. The strains SB, GN and F15 are derived from Zymaflore VL1, Actiflore BO213, Zymaflore F15 (Laffort, Bordeaux, France), respectively, while M2 is derived from Oenoferm M2 (Lallemand, Blagnac, France). The remaining 31 strains used are commercial starters obtained from different companies. To avoid any conflict of interest there were encoded C1 to C31 and are available and deposited on the CRB collection of ISVV, S1Table. Yeasts were propagated on YPD (Yeast extract 1 % Peptone 1 % Dextrose 2 %) supplemented with agar (2%) when required. The strains were long-term stored in YPD with 50 % of glycerol at -80 °C.

### Grape musts and vessels used and fermentation monitoring

The five grape musts used, i.e. Merlot 2014 (M14), Merlot 2015 (M15), Cabernet Sauvignon 2014 (CS14), Sauvignon Blanc 2014 (SB14) and Sauvignon Blanc 2015 (SB15), were provided by *Vignobles Ducourt* (Ladaux, France) and stored at -20 °C. Before fermentation, grape musts were sterilized by membrane filtration (cellulose acetate 0.45 μm Sartorius Stedim Biotech, Aubagne, France). Their main enological characteristics are given in Table 1. Sugar content, assimilable nitrogen, pH, total and free SO_2_ have been assayed by the enological analysis laboratory (SARCO, Floirac, France). Malic acid has been assayed by enzymatic essay as described in the enzymatic assay section. Initial active SO_2_ concentration was estimated using the protocol given at http://www.vignevin-sudouest.com/services-professionnels/formulaires-calcul/so2-actif.php. Input parameters used: pH and free SO_2_ concentration of the grape must, fermentation temperature (24 °C), and 0.1 % of alcohol by volume to simulate the beginning of the fermentation.

**Table 1:** Grape musts composition

| Grape must | Code | Sugar<br>content<br>(g.L<sup>-1</sup>) | Assimilable<br>Nitrogen (mg N.L<sup>-1</sup>) | Malic acid<br>(g.L<sup>-1</sup>) | pH | total SO<sub>2</sub><br>(mg.L<sup>-1</sup>) | free SO<sub>2</sub><br>(mg.L<sup>-1</sup>) | active SO<sub>2</sub><br>(mg.L<sup>-1</sup>) |
| --- | --- | --- | --- | --- | --- | --- | --- | --- |
| Sauvignon Blanc 2014 | SB14 | 194 | 157 | 5.6 | 3.19 | 34 | 7 | 0.32 |
| Sauvignon Blanc 2015 | SB15 | 203 | 158 | 2.9 | 3.25 | 67 | 23 | 0.91 |
| Merlot 2014 | M14 | 207 | 111 | 2.1 | 3.58 | 37 | 29 | 0.54 |
| Merlot 2015 | M15 | 219 | 99 | 1.9 | 3.53 | 46 | 33 | 0.68 |
| Cabernet Sauvignon 2015 | CS15 | 220 | 132 | 2.4 | 3.57 | 35 | 25 | 0.47 |

To carry out the fermentations, 10 mL screwed vials (Fisher Scientific, Hampton, New Hampshire, USA) were used in order to ferment 3 mL or 5 mL of grape must in a standardized way. The Screwed Vials (here after named SV) were tightly closed with 18mm screw cap-magnetic- 3mm HT silicone/PTFE (Fisher Scientific, Hampton, New Hampshire, USA). Hypodermic needles (G 26 – 0.45 x 13 mm, Terumo, Shibuya, Tokyo, Japan) were inserted into the septum for CO_2_ release.

Fermentations were initiated by inoculating 2.10^6^ viable cell.mL^-1^ of 24h-liquid culture (YPD) carried out in 1 mL deepwell microplates (Fisher Scientific, Hampton, New Hampshire, USA). The concentration of viable cells was estimated by flow cytometry using a Cell Lab Quanta apparatus (Beckman Coulter, Brea, California, USA) according to the method described by Zimmer *et al*. [8].

The fermentation temperature was maintained at 24°C by an incubator (Binder GmbH, Tuttlingen, Germany). When specified, the SV were shaken at 175 rpm during the overall fermentation using an orbital shaker (SSL1, Stuart, Vernon Hills, Illinois, USA). In order to compare this new vessel type with already published conditions, 125 mL-glass bioreactors (GB) were also used according to the specification described by da Silva *et al*. [29].

The fermentation kinetics were estimated by monitoring regularly the weight loss caused by CO_2_ release using a precision balance (AB104, Mettler Toledo, Greifensee, Switzerland). Theoretical maximum CO_2_ release (*tCO*_*2*_*max*) was calculated according to the formula: 0.482*[Sugar] [29], where [Sugar] is the sugar concentration (g.L^-1^) of the must. The amount of CO_2_ released according to time was modeled by local polynomial regression fitting with the R-loess function setting the span parameter to 0.45. Six kinetic parameters were extracted from the model:

- *lp* (h): lag phase time observed before to release the first 2 g.L^-1^ of CO_2_;
- *t35*, *t50* and *t80* (h): time to release 35, 50 and 80 % of the tCO_2_max after subtracting *lp*;
- *V50_80* (g.L^-1^.h^-1^): average CO_2_ production rate between 50 % and 80 % of *tCO2maX*;
- *CO*_*2*_*max*: maximal amount of CO_2_ released (g.L^-1^).

### Enzymatic assays

At the end of the fermentation, a sample volume of 800 μL was stored at -20 °C and analyzed at the metabolomics platform of Bordeaux by semi-automatized enzymatic assays (http://metabolome.cgfb.u-bordeaux.fr/). The concentrations of the following organic metabolites were measured: acetic acid, glycerol, malic acid, pyruvate, acetaldehyde and total SO_2_ using the respective enzymatic kits: K-ACETGK, K-GCROLGK, K-LMAL-116A, K-PYRUV, K-ACHYD, K-TSULPH (Megazyme, Bray, Ireland) following the instructions of the manufacturer. Dilution level and volume of sample used are described in S2 Table. Glucose and fructose were assayed by using the enzymatic method described by Stitt et al. [30], however in the presented data, all the fermentations were completed containing less than 1.5 g.L^-1^ of residual sugars.

### Apolar esters analysis

Samples were analyzed after thawing. Concentration of 32 esters (ethyl fatty acid esters, acetates of higher alcohol, ethyl branched acid esters, isoamyl esters of fatty acid, methyl fatty acid esters, cinnamates and minor esters) (S3 Table). Concentration was determined using a head space solid phase microextraction (HS-SPME) followed by gas chromatography–mass spectrometry (GC–MS) as described by Antalick *et al.* [31].

### Dissolved oxygen measurement

To control the initial oxygen concentration, oxygen was removed by bubbling nitrogen inside SV for 20 min. Non-intrusive measurement of the concentration of dissolved oxygen in the grape juice was done by using NomaSense O2 P300 sensor (Nomacorc, Narbonnes, France) bonded on the inner surface of the SV.

### Statistical analyses

All the statistical and graphical analyses were carried out using R software [32]. The variation of each trait was estimated by the analysis of variance (ANOVA) using the *aovp* function of the *lmPerm* package in which significance of the results was evaluated by permutation tests instead of normal theory tests. Tukey's honest significant difference test was used on *aovp* results to determine which group of means differ significantly using the *HSD.test* function (*agricolae* package) [33].

The LM1 model estimated the effect of strain, of grape must of micro-oxygenation of the strain-by-must interaction and of the strain-by-micro-oxygenation interaction on fermentation traits according to the following formula:

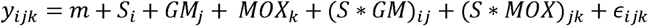

where y_ijk_ was the value of the trait for strain *i* (*i* = 1, …, 4) in grape must *j* (*j* = 1, 2) and with micro-oxygenation level *k* (*k* = 1, 2), m was the overall mean, *S*_*i*_ was the strain effect, *GM*_*j*_ the grape must effect, *MOX*_*k*_ the micro-oxygenation effect, *(S * GM)_ij_* was the interaction effect between strain and grape must, *(S * MOX)_jk_* was the interaction effect between strain and micro-oxygenation level and *∊*_*ijk*_ the residual error. Correlations between traits were computed with the Spearman method using the *cor* function and the significance of the results was assessed by the *cor.test* function at 0.95 of confidence level. Results were displayed with the *corrplot* function (*corrplot* package).

Principal Component Analysis (PCA) was calculated using the *ade4* package and heatmaps were generated with the *heatmap.2* function. When necessary non-parametric comparison of samples were carried out using the Wilcoxon-Mann-Withney test (α = 0.05).

## Results

### Optimization of the fermentation protocol in screw capped vials

The first aim of this study was to develop a fermentation method for measuring in a reliable manner numerous strains in a small volume (<10 mL). We used 10 mL-screwed vials (SV) filled with 3 or 5 mL of grape must. Their small and standard size can be conveniently exploited to run in parallel more than 300 fermentations at the same time in a small space (S1 Fig). In preliminary experiments (not shown), we observed that the volume of grape juice used influences the success of the fermentation. To evaluate this effect on enological parameters, the fermentation behavior of four yeast strains (M2, F15, SB, GN) was evaluated in the SB14 grape must in 6 replicates. Three conditions were tested: 3 mL with shaking (Sk.3_SV), 5 mL with shaking (Sk.5_SV) and 5 mL without shaking (noSk.5_SV). In order to validate the SV, the same juice was also fermented in 125 mL glass-bioreactors (Sk.125_GB) that had been previously used for measuring the fermentation behavior of numerous *Saccharomyces* strains and hybrids [29]. For all assays, fermentations were completed (no residual sugars detected); the overall results are given in the S4 Table for the 12 parameters measured for each strain in the 4 assays.

To compare the reliability of trait values, the coefficient of variation (CV %, for 6 replicates) was computed for each strain and the average CV was shown in Table 2. The fermentation kinetic traits are very reliable confirming the efficiency of weight loss measurement for monitoring ongoing alcoholic fermentations [34], even in very small volumes (Fig 1, panel A). For some metabolic traits, high CVs (>25 %) were measured showing that some conditions are not reliable enough. This is the case for acetaldehyde, pyruvate or acetic acid for which the CVs are particularly high in shaken conditions. The Sk.3_SV trial was the less reliable and the cumulated CV for metabolic compounds is much higher than for the other 3 conditions (Fig 1, panel B). In this condition, the kinetics parameters are also less reproducible (CV>10 %). In contrast, noSk.5_SV offers the most reliable condition for both metabolic compounds and kinetic parameters. Except for the lag phase, the Sk.5_SV condition had an intermediate reliability level, similar to the 125 mL glass-bioreactors used here as a control.

**Table 2:** Average coefficient of variation for the different traits

| Condition | CO<sub>2</sub>max | Lp | t35 | t50 | t80 | V50_80 | SO<sub>2</sub> | Acetic acid | Glycerol | Malic acid | Pyruvate | Acetaldehyde |
| --- | --- | --- | --- | --- | --- | --- | --- | --- | --- | --- | --- | --- |
| noSk.5_SV | 2.0 | 11.5 | 6.8 | 7.4 | 7.3 | 7.3 | 9.1 | 11.3 | 6.4 | 5.4 | 17.8 | 17.4 |
| Sk.125_GB | 2.0 | 9.8 | 6.4 | 5.0 | 5.3 | 6.8 | 9.8 | 13.8 | 11.8 | 19.0 | 45.8 | 32.1 |
| Sk.3_SV | 0.8 | 21.4 | 8.6 | 8.4 | 9.2 | 13.6 | 6.0 | 61.1 | 17.0 | 14.8 | 63.2 | 46.2 |
| Sk.5_SV | 2.3 | 41.2 | 8.0 | 7.3 | 6.1 | 6.7 | 6.2 | 25.0 | 12.1 | 19.1 | 27.4 | 26.2 |

The data presented are the average coefficients of variation (CV in %) calculated from the CV values obtained for each strain with 6 replicates

**Fig 1.**
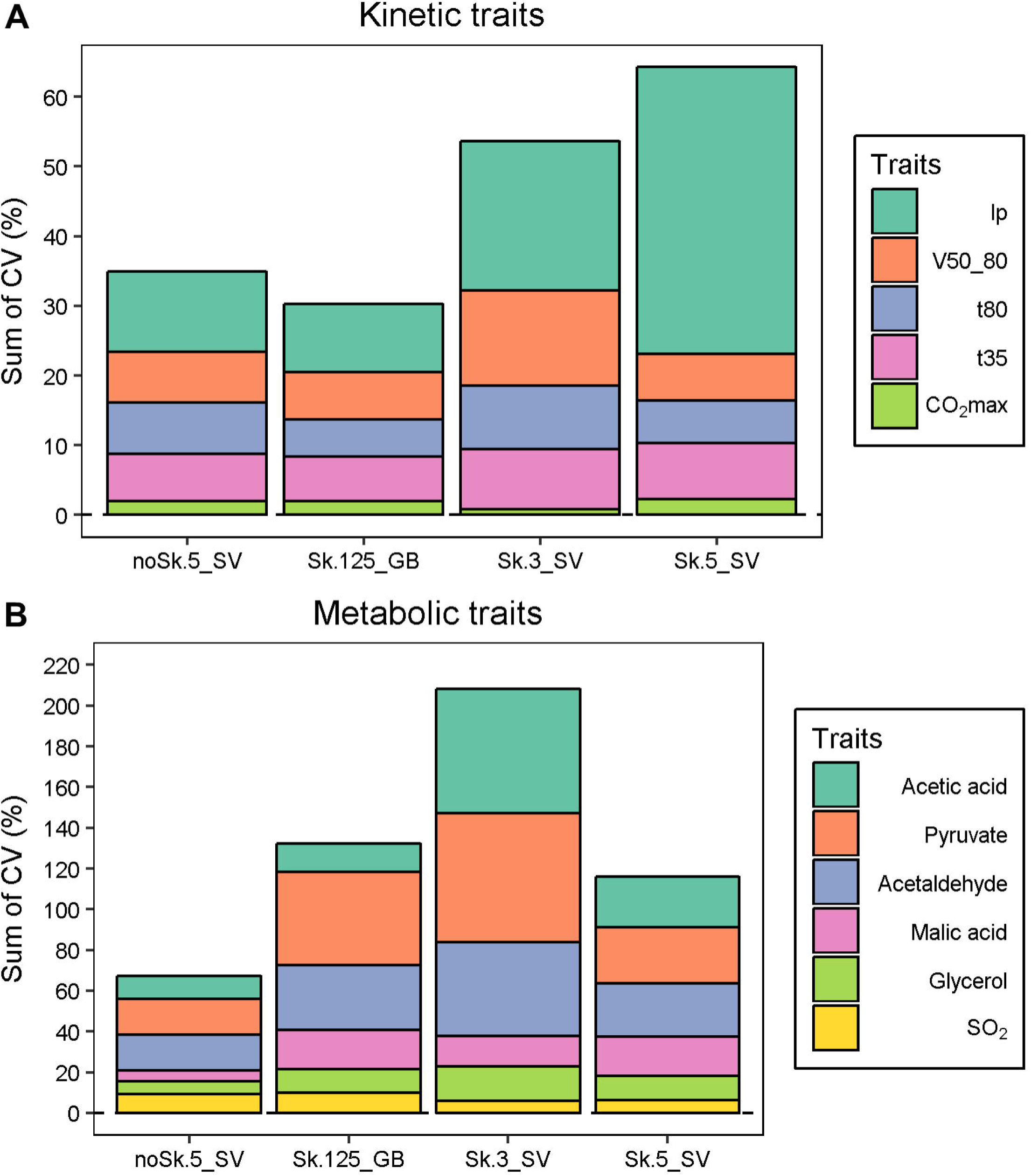
Trait measurement reliability for both kinetics and metabolite concentrations according to vessel modalities. The average CV for each trait was calculated from the CV values obtained for each strain (M2, F15, SB, GN) with 6 replicates. Panel A. The bar chart presents the cumulated CV for each kinetic parameter, the stacking is ordered from the least variable (*CO*_*2*_*max*) to the most variable (*lp*) trait. Panel B. The bar chart presents the cumulated CV for each metabolic end-product, the stacking is ordered from the least variable (*SO*_*2*_) to the most variable (*Acetic acid*).

Despite important changes according to the conditions, the overall differences between the four strains were maintained and the genetic differences within the strains were broadly conserved (see below). Strikingly, the shaking conditions impacted the fermentation kinetics for all the strain. This is illustrated for example with the CO_2_ kinetics of the GN strain, Fig 2, panel A. The CO_2_ production rate was dramatically impacted by shaking, which significantly reduced (by around 20 %) the *t50* and *t80* (Wilcoxon test α = 0.01). In contrast, the fermentation volume (3, 5 and 125 mL) did not affect the fermentation kinetics in shaken conditions, suggesting that scaling down in SV did not influence the fermentation behavior of yeast cell. The metabolic end-products were also affected by the shaking conditions, as shown in Fig 2, panel B for glycerol. As observed for kinetic parameters, the fermentation volume had a minor impact on the primary metabolites composition (such as glycerol) whereas shaking appeared as the main source of phenotypic variation. This result, observed for all strains, could be due to the higher oxidative conditions met in shaken cultures.

**Fig 2.**
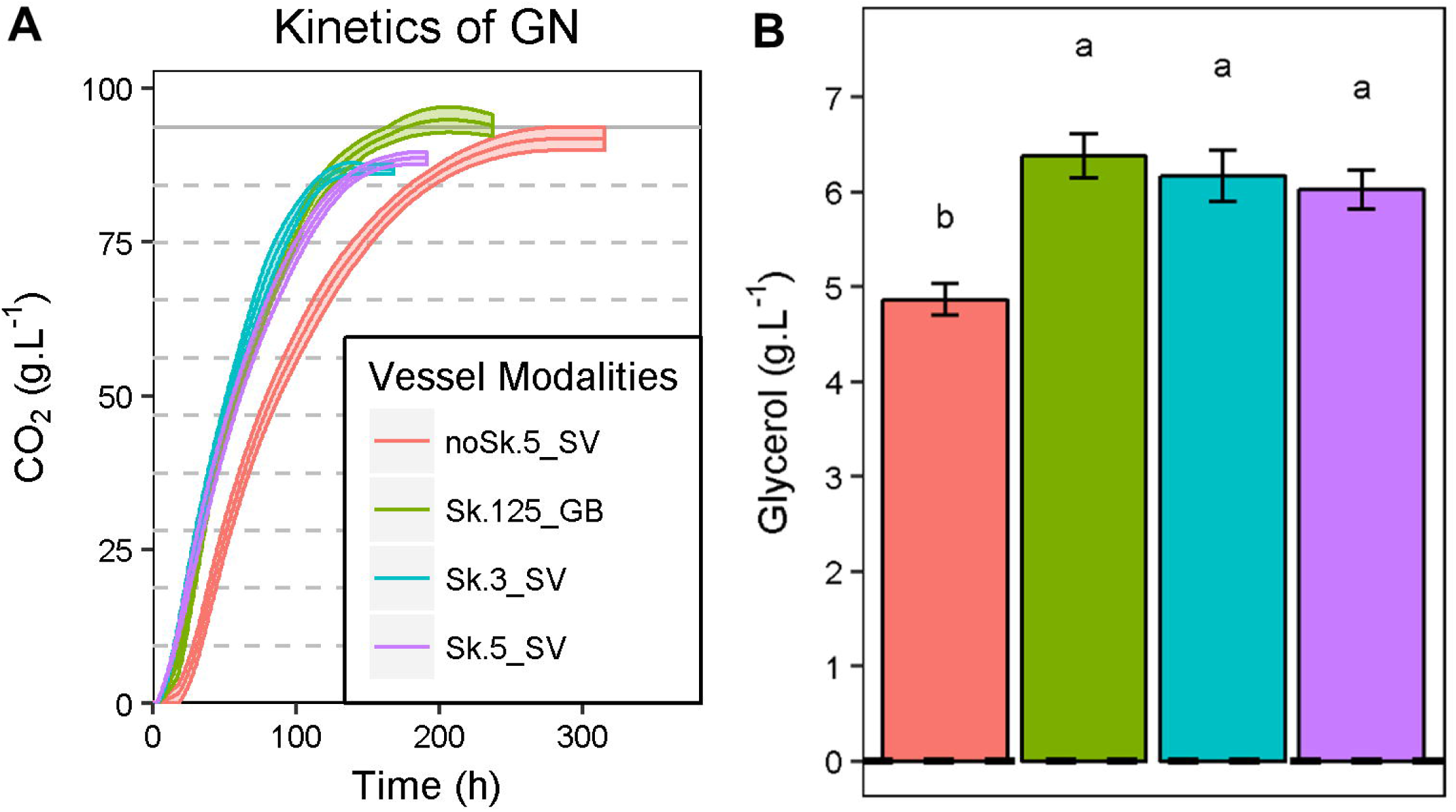
Impact of agitation on fermentation kinetics and metabolic compounds according to the vessel modalities. Panel A. CO_2_ production kinetics of the GN strain fermenting SB14 grape must in four vessel modalities (Sk.3_SV, Sk.5_SV, noSk.5_SV, Sk.125_GB). The lines are the average CO_2_ produced for 6 replicates; the shaded areas represent the standard error. Panel B. Glycerol production of GN strain according the vessel modalities. The values shown are the means of 6 replicates and the error bars represent standard error.

A second experiment was performed in 5 mL SV as they represent the most reproducible conditions for measuring all the traits investigated (Fig 3). The micro-oxygenation effect was estimated by comparing modalities with or without shaking during the fermentation. The O_2_ concentration was monitored during 20 hours in non-inoculated SB14 grape juice degassed by nitrogen bubbling. During this period, corresponding to the fermentation lag phase, oxygen can be efficiently transferred since CO_2_ stripping is not active. Although this measurement did not correspond to real conditions since no yeast cells were present, the effect of agitation on the oxygen transfer could be estimated. Indeed, when yeast cells are present, all the dissolved oxygen is consumed in less than 20 hours due to the strong reductive conditions generated by yeast biomass (data not shown). In the shaken condition, the grape juice was immediately enriched with dissolved oxygen that reached a concentration of 3.7 mg.L^-1^ after 20 h (Fig 3, panel A). In contrast, without shaking, there was only 2.4 mg.L^-1^ of dissolved oxygen after 20 h. A maximum difference in oxygenation rate was found after 3 hours of incubation (Fig 3, panel B). Although the total amount of oxygen transferred during the overall fermentation cannot be measured, these data suggest that agitation in 5 mL-SV significantly impacts the micro-oxygenation level. These small, but significant differences may explain the kinetic and metabolic differences described in Fig 2.

**Fig 3.**
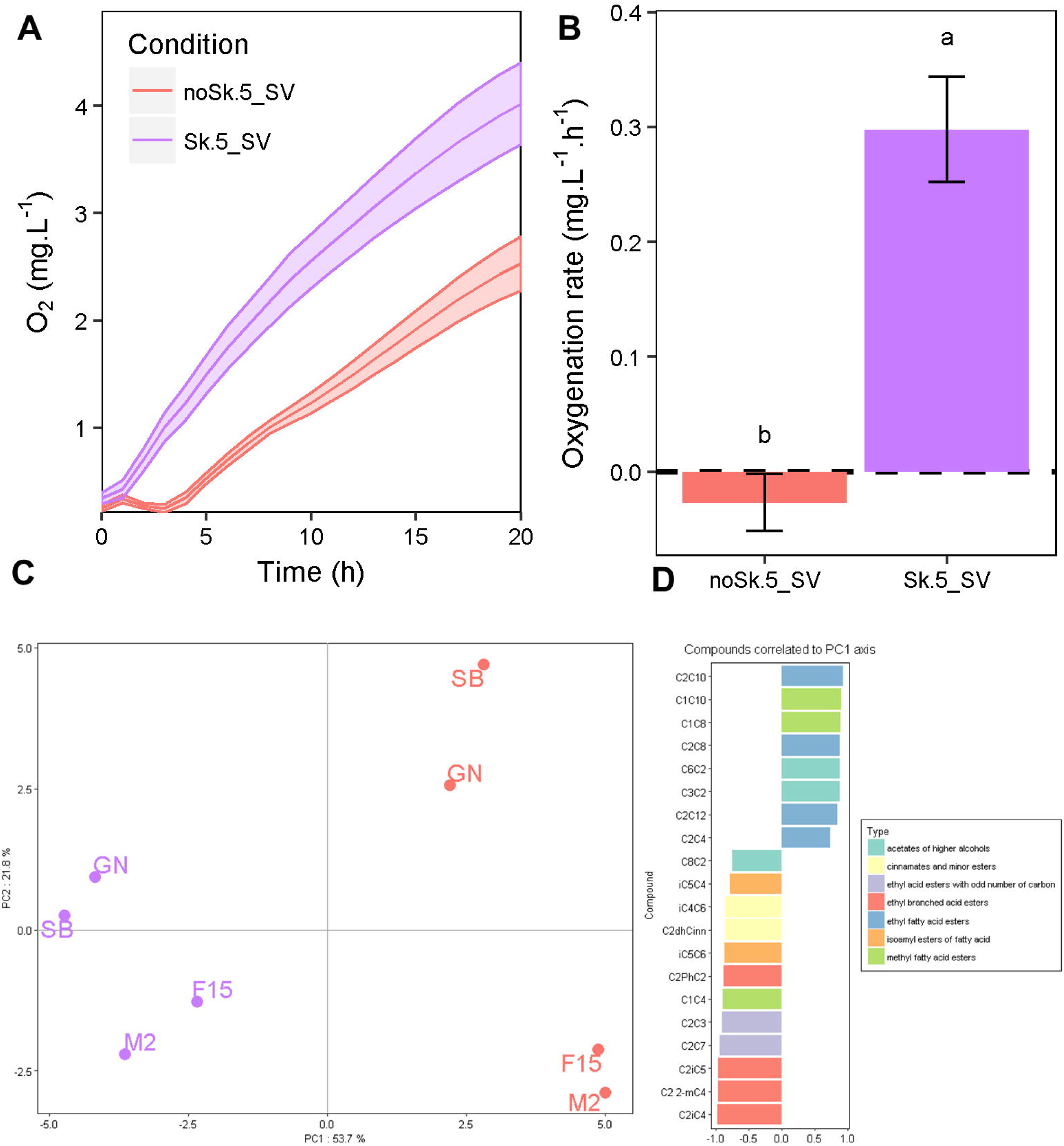
Measure and effect of micro-oxygenation in 5 mL SV. Panel A. Kinetics of dissolved oxygen concentration in SB14 grape must. The kinetic curves represent the mean of 6 replicates and the shadows around the lines illustrated the standard errors. Panel B. Concentration of the dissolved oxygen in SB14 after 4 hours. The data shown are the means of 6 replicates and the error bars represent the standard deviations. Different letters indicate significant differences between groups (Tukey's honest significant difference test, significance level, α = 0.05). Panel C. PCA performed for the 32 esters measured. Each point represents one of the four the strains in noSk.5_SV or in Sk.5_SV. Panel D. Correlation of the variables to the PCA1 axis. The variables that were significantly correlated to the first axis of the PCA were shown (α = 0.05), the bar plot indicated the pval of the correlation (Pearson's product moment correlation coefficient).

In order to have a broader idea of the impact of micro-oxygenation on secondary metabolism, we next measured the production of volatile compounds. At the end of the alcoholic fermentation, the headspace volume of SV was analyzed using a targeted GC-MS analysis. 32 esters were quantified for the fours strains in shaken or not conditions (S5 Table). A Principal Component Analysis (PCA) (75.5 % of total variance for axes 1 and 2) was carried out for exploring this multivariate dataset (Fig 3, panel C). The first component clearly discriminates shaken from non-shaken conditions while the second axis mainly discriminates strains. Indeed, the production of esters was greatly impacted by shaking. Up to 27 of the 32 esters were significantly impacted (ANOVA, pval<0.05), 14 with a decreased and 13 with an increased production in the shaken condition (S5 Table). The compounds, for which shaking decreased their production, were mainly acetates of higher alcohols, methyl and ethyl fatty acid esters while those for which the production was increased were mainly ethyl branched acid esters, ethyl acid esters with odd carbon numbers, cinnamates and minor esters. The proportion of PhC2C2 to C2PhC2 was 6 fold decreased in shaken condition (S2 Fig). This could be caused by a higher oxygenation of the media.

### Assessment of genetics x environmental effects

In order to demonstrate the efficiency of our SV fermentation setup, we explored phenotypic response of strains to relevant environment parameters in enology. On the basis of the results shown in Fig 2-3, shaken fermentations could be considered as micro-oxygenated modalities transferring moderate amounts (2-4 mg.L^-1^ per day) of oxygen in a reproducible way. The possibility to control oxygenation in small volumes is an opportunity to study the reaction of yeast strains against this technological parameter which has a significant impact on winemaking [35–37]. Assuming this statement, a second experiment was carried out in 5-SV, by fermenting the two grape juices SB14 and M15 with four strains (M2, GN, F15 and SB) and with or without shaking. This set of 160 fermentations (S6 Table) ran at the same time allowed to estimate the effects of three main factors: (i) strain, (ii) micro-oxygenation, and (iii) grape must. The proposed model for the analysis of variance also estimated the primary interaction within strain and grape must or micro-oxygenation (model LM1 described in material and methods). Thanks to the small volume used, 10 biological replicates were carried out for each strain and condition, thus increasing the statistical power of the analysis. For most of the traits, the phenotypic variance was first explained by the grape juice type, then by the yeast strain used (Table 3). The effect of micro-oxygenation mainly influenced kinetic parameters (*t50*, *t80*) and metabolic end-product such as *SO*_*2*_ and *Glycerol.* For this last trait, the micro-oxygenation increased the production by 15 % (Fig 4, panel A) for all the strain, as previously reported by others [38–40]. Few strain *x* environment interactions were detected and accounted only for a small part of the total variance explained. The most striking interaction pertained to the lag phase duration (*lp*) being differentially affected by the micro-oxygenation and the grape must, respectively. The panel B of Fig 4 shows that the strains SB and M2 had a longer lag phase in the SB14 grape must than in M15 (+ 6 h). Moreover, shaking resulted in a reduced lag phase for M2 in the SB14 grape must. In contrast, F15 and GN were not affected neither by the grape must nor by the agitation. In the same way, the *acetic acid* production of GN showed a complex GxE interaction (Fig 4, panel C). Globally, as previously described [41], micro-oxygenated conditions tended to reduce the production of this compound, which is undesirable in enology. Interestingly, in the M15 grape must, GN showed the lowest acetic acid production even in a non-agitated fermentation, suggesting that it is an interesting lower producer whatever the conditions. This second experiment confirms the reliability of SV for assessing wine fermentation traits in various environmental conditions and paves the way for larger phenotypic investigations.

**Table 3:** Analysis of variance for the 11 phenotypes with 4 strains, 2 musts and 2 micro-oxygenation conditions

|  | CO<sub>2</sub>max | lp | t35 | t50 | t80 | V50_80 | SO<sub>2</sub> | Acetic acid | Malic acid | Pyruvate | Glycerol |
| --- | --- | --- | --- | --- | --- | --- | --- | --- | --- | --- | --- |
| **Must** | 37.9 *** | 15.8 *** | 38.6 *** | 36.8 *** | 35.2 *** | 21.5 *** | 15.4 *** | 10.7 *** | 81.2 *** | 4.9 ** | 0.3 |
| **Strain** | 2.3 . | 41.5 *** | 10.3 *** | 16.7 *** | 27.3 *** | 43.8 *** | 4.3 *** | 3.7 * | 8.1 *** | 9 *** | 17.2 *** |
| **Micro-Oxygenation** | 7.4 *** | 2.8 *** | 37.2 *** | 32.1 *** | 22 *** | 20.2 *** | 40.3 *** | 39.1 *** | 0 | 5.2 *** | 49.4 *** |
| **Strain:Must** | 0.3 | 10 *** | 2.1 *** | 2.5 *** | 2.7 *** | 0.5 | 2.8 ** | 0.4 | 0.2 | 6.8 *** | 0.1 |
| Strain:Micro-Oxygenation | 0.6 | 3.7 *** | 0.7 * | 0.3 | 1.6 *** | 0.1 | 1.8 . | 2.6 * | 1.1 *** | 4 . | 1.8 * |
| Residuals | 51.4 | 26.2 | 11.1 | 11.6 | 11.3 | 13.9 | 35.5 | 43.5 | 9.4 | 70 | 31.3 |

Percentage of variance explained by the LM1 model. Significance codes: $pval < 0.001 = ***$ ,
$pval < 0.01 = **$ , $pval < 0.05 = *$ , $pval < 0.1 = .$

**Fig 4.**
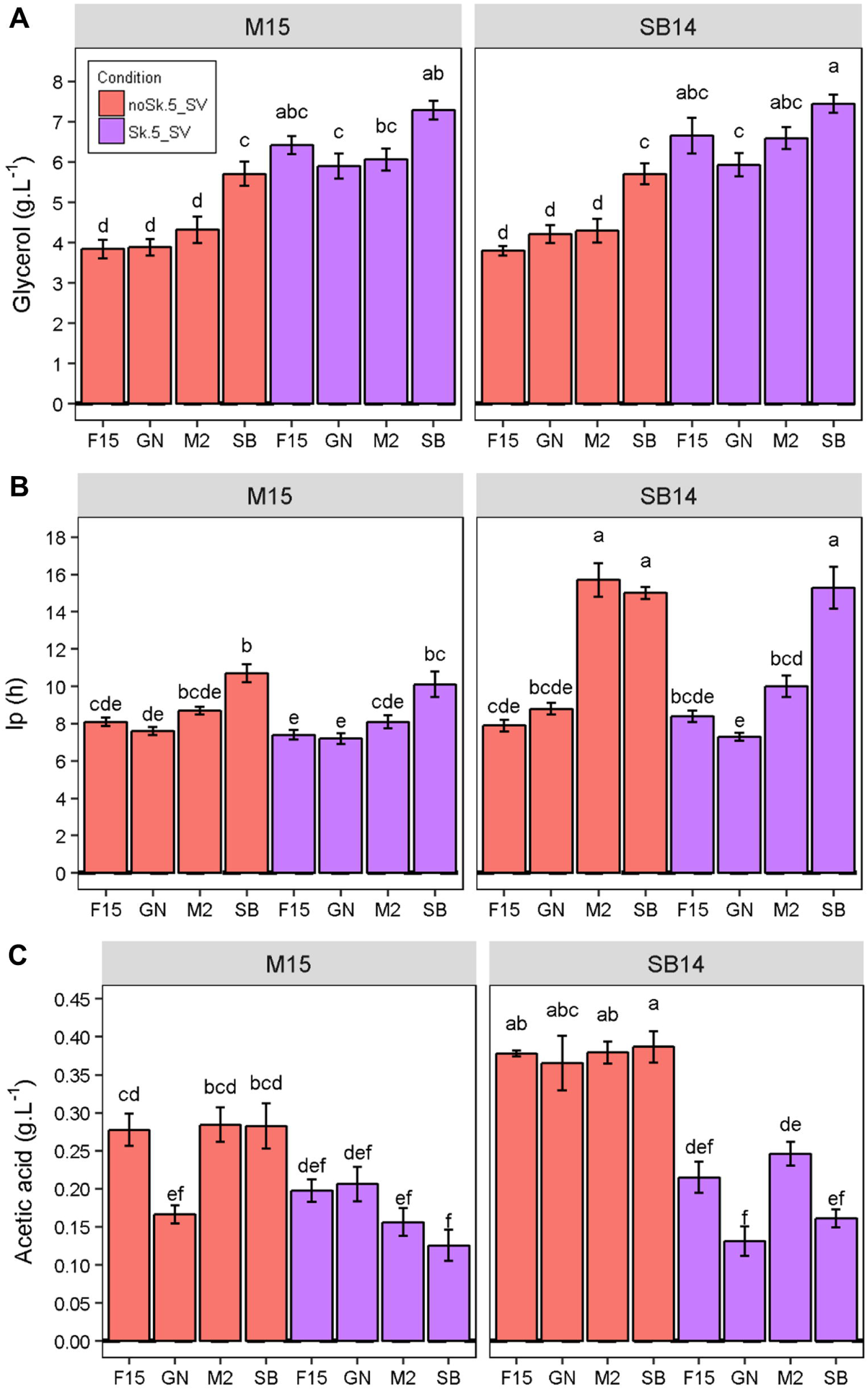
Effect of micro-oxygenation level and grape must on technological properties of wine yeast strains. The data shown are the mean of 10 replicates, the error bars representing the standard error. Different letters indicate significant differences between groups (Tukey's honest significant difference test, significance level, α = 0.05). Panel A. *Glycerol* (g.L^-1^) according to strain and fermentation conditions. Panel B. *lp* (h) according to strain and fermentation conditions. Panel C. *Acetic acid* (g.L^-1^) according to strain and fermentation conditions.

### Evaluation of technological properties of 35 wine yeast strains in 5 grape juices

The SV fermentation setup coupled with robotic assisted enzymatic assays offers the opportunity to measure in parallel the fermentation behavior of numerous strains in various conditions. As a matter of proof, we evaluated in a unique experiment the fermentation properties (kinetics and end by-products) of 35 strains in 5 grape juices and two repetitions (350 SV) without shaking. In this experiment, we used three red grape musts (M14, M15 and CS14) and two white grape musts (SB14 and SB15) from the Bordeaux area. As all the fermentations were completed (less than 1.5 g.L^-1^ of residual sugars), the final concentrations of glucose and fructose were very low and thus removed from the data (not shown). Acetaldehyde concentrations were also removed, as they were very low in red wines and thus impacted data normality (not shown). The measurement of the 11 quantitative variables for 175 modalities is given in S7 Table. For all the traits analyzed, except pyruvate, the average CV per trait (n = 175) was less than 18 %.

A PCA (58 % of total variance for axes 1 and 2) was carried out for exploring this large dataset. The first component (42 % of total variance) clearly discriminates red and white juices and was correlated with *Malic acid*, *SO*_*2*_, *Acetic acid* concentrations and kinetic parameters (*t50*, *t80*, *V50_80*) (Fig 5, panel A). Indeed thewhite grape juices used were more acidic and more sulphited than red ones. The second axis (16 % of total variance); mainly discriminates the CS15 must from the others by its higher production of *glycerol* and *CO*_*2*_*max*. These results are consistent with the biochemical composition of grape juices. Moreover the CS15 juice contained 20 g.L^-1^ more sugar than the other grape musts.

**Fig 5.**
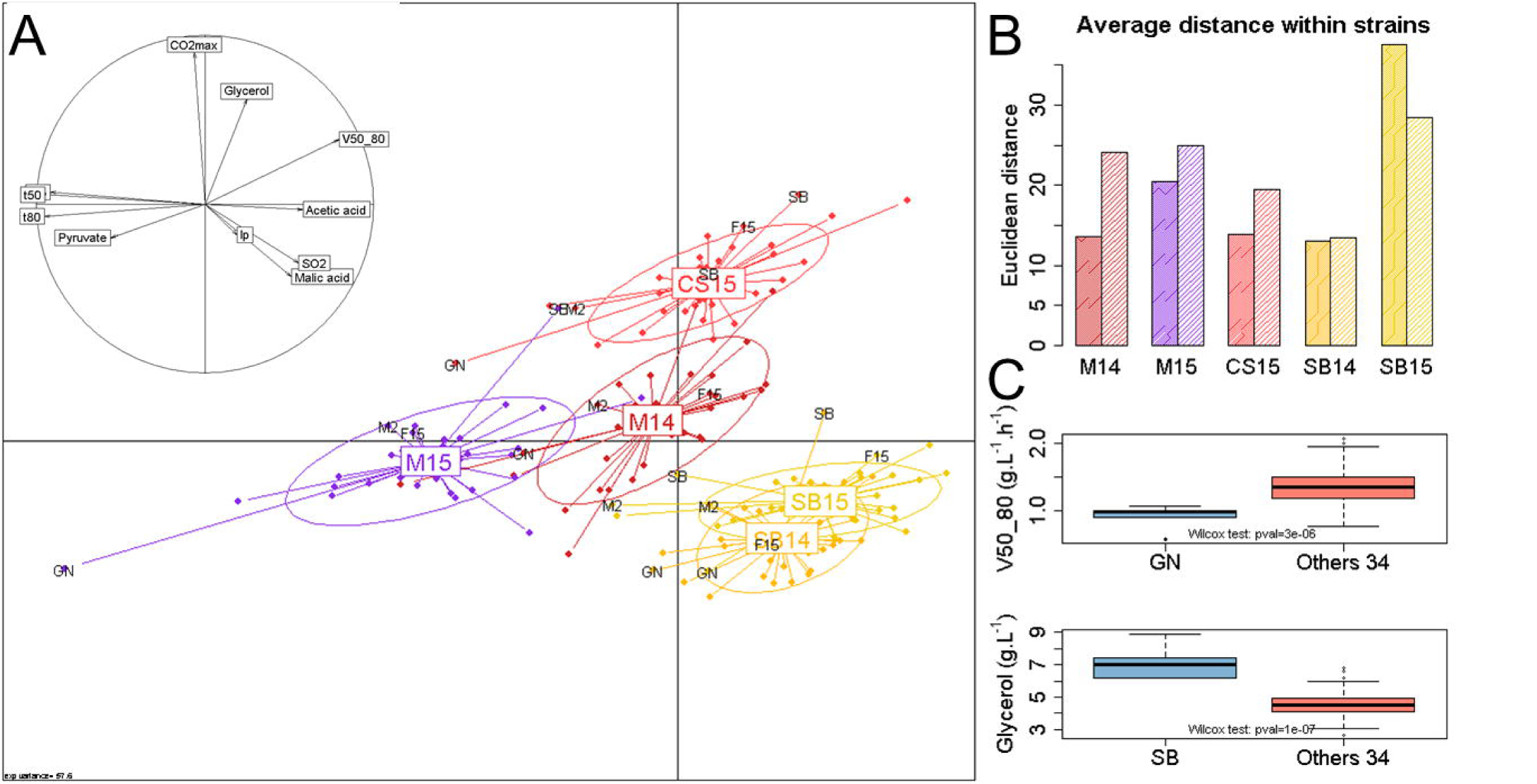
PCA of winemaking properties of 35 strains in 5 grape juices. Panel A. The first two axes of the PCA performed from the average of two replicates for 11 phenotypes measured in the 5 grape juices and 35 strains. Axes 1 and 2 explain 41.8 % and 15.8 % of total variation, respectively. Each point represents the fermentation of one strain and is colored according to the grape juice used. Points are connected to their group gravity centers that are labeled with the grape juice name M14, M15, SB14, SB15, CS14. Ellipses diameter corresponds to the standard deviations of the projection coordinates on the axes. The correlation circle indicates the correlation of the variables for axes 1 and 2. Panel B. Euclidian distances within all the strains for each grape must. The bar plot represents the Euclidian distances within the 35 strains according to kinetics (high density colored bar) and metabolic parameters (low density colored bar) for each grape juice. Panel C. Comparison of the trait value of GN and SB respect to the 34 others strains for V50_80 and the glycerol produced, respectively. A boxplot was generated from the 10 phenotypic values measured in the 5 grape juices with two replicates for GN and SB, and from the 340 values of the 34 other strains. Significant differences were estimated by applying the Wilcoxon-Mann-Withney test (α = 0.05).

The PCA also illustrates the phenotypic variability of the 35 industrial strains tested. Globally, the analysis showed that some grape musts are more suitable than others for between strains. Indeed, the projected cloud of the 35 strains in SB14 is more compact than in M15. In order to evaluate the discriminating properties of each grape must, we computed the average Euclidian distance within all the strains for both kinetic and metabolic parameters and according to the grape must. The panel B of Fig 5 summarizes the phenotypic distance observed within each grape must and parameter class. For example, SB15 emphasized strain discrepancy for kinetic traits and metabolic end-products. To better visualize particular strain properties, the positions of the four strains SB, GN, M2 and F15 were labeled on the projection. These strains have some phenotypic specificities; for example SB and GN are often more distant from the remaining set of commercial strains than M2 and F15. This is in particular due to the high *glycerol* production of SB and the slow fermentation rate (*V50_80*) of GN in all the conditions tested (Fig 5, panel C).

As shown on the PCA, the nature of the juice strongly impacted the phenotypic values. In order to overcome this effect and perform more accurate comparative analyses between strains, we normalized the response of each strain according to the grape juice (S8 Table). First, the relations between the 11 traits were investigated by using the average of normalized values of each strain for the five conditions. A correlation matrix with non-parametric tests was computed with the 35 strain values in order to observe phenotype-phenotype relations (Fig 6, panel A). Obvious correlations between kinetic traits were found confirming that the strains that rapidly reached 35 % of the fermentation (lowest *t35* values) had also low *t80* values (S3 Fig). Interestingly, we detected less trivial correlations suggesting metabolic link. For example, a correlation between kinetic parameters and *Malic acid* was found (Fig 6, panel B). The strains with the fastest fermentation rates were also the ones that consumed the most of malic acid. This link has already been reported [42] and could be explained by a greater deacidification capacity for strains that consume more malic acid, resulting in easier fermentation. Negative correlations were found between kinetic parameters (*t35*, *t50*, *t80*) and *SO*_2_. These negative relations could be explained by the toxic effect of SO_2_ that reduces yeast growth [43,44] and may indirectly impact the fermentation activity. Other correlations were found for *lp* with *V50_80* (Fig 6, panel C) and *glycerol* and will be discussed further.

**Fig 6.**
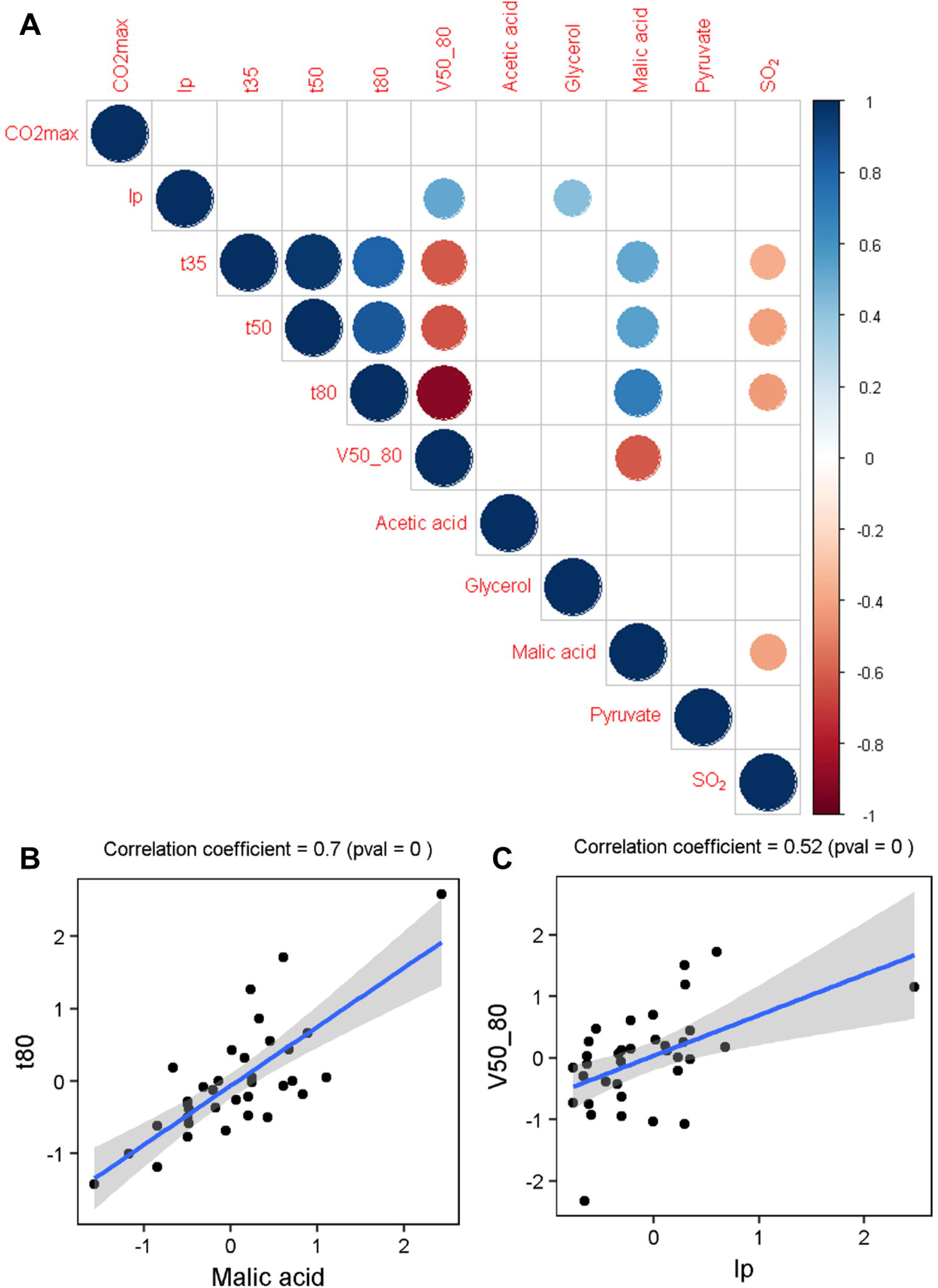
Correlation between traits. Panel A. A correlation matrix is shown. The size and the colour of the circles correspond to the correlation coefficients calculated by the Spearman method. Only significant correlations are shown (confidence = 0.95). Panel B and C. Two examples of scatter plots showing correlation of *t80* with *Malic acid* and *V50_80* with *lp*. Each dot represents the average phenotypic values of a strain across the 5 grape musts from the normalized dataset. The blue line represents the linear regression line and the shaded area represents the confidence interval of the regression (0.95).

The normalized dataset was also used for evaluating the performance of the strains. The rank of each strain with respect to the others was calculated and can be visualized on a heatmap plot (Fig 7). As each column of the heatmap plot represents a rank value (1 to 35), each trait has the same weight in the clustering. Because most of the kinetic parameters are strongly correlated (Fig 6, panel A), only three of them (poorly correlated) were included in the analysis (CO_2_max, *lp* and *V50_80*). The intensive green tones indicate lowest ranks while intensive red tones indicated the highest ranks for each parameter. For example, the commercial strains C11, C4 and C18 were among the fastest strains and consumed more malic acid than the others. Rapid identification of strains having outlier levels compared to a representative commercial set can be made with this figure. For example, the strains C6, C17 and C20 produced high quantities of acetic acid while the strains C5, C8 and C16 released an important quantity of SO_2_ at the end of the alcoholic fermentation. As displayed by the dendrogram on the left of the heatmap, a hierarchical clustering ordered the strains according to their overall profiles. Four main groups were computed. The group A contained slow fermenting strains, which leave high amounts of *malic acid* at the end of the fermentation and produce low *SO*_*2*_. Group B contained strains with the shortest *lp.* Moreover, most of the strains of this group had a slow fermentation rate, produced low amounts of *glycerol* and released high level of *SO*_*2*_. The strains of group C were the fastest fermenting ones, produced more *glycerol* and *SO*_*2*_ than the average. This group also consumed more *malic acid*. Finally, the strains of group D fermented rapidly but in contrast with those of group C they produced low amounts of *glycerol* and *SO*_*2*_.

**Fig 7.**
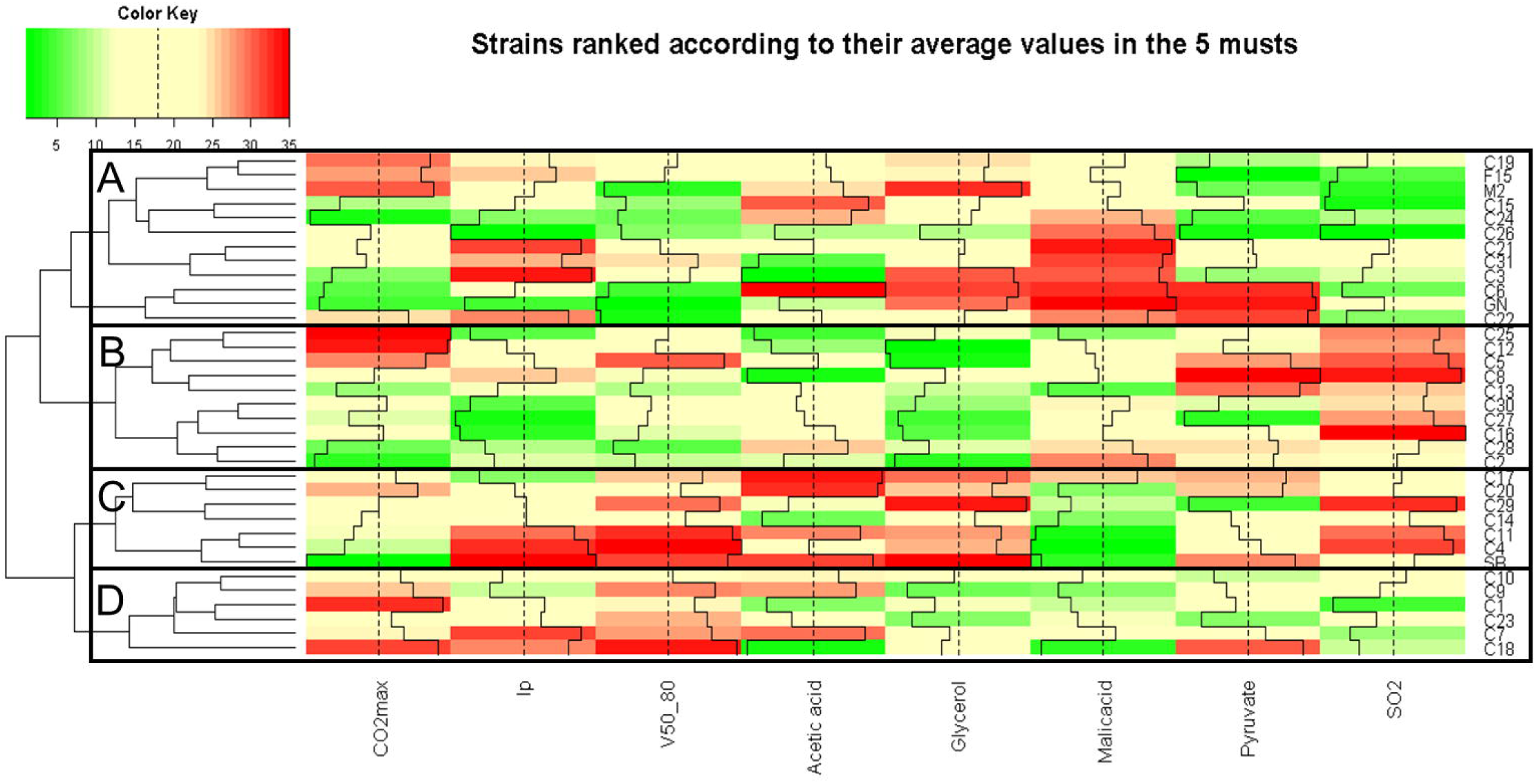
Relative ranking of 35 strains in 5 grape juices. Ascending order ranked of the average phenotypic values of each strain across the 5 grape juices. Only a subset of the representative phenotypes is represented here. A color palette shows each rank from green (lowest ranks) to red (highest rank) as displayed by the color key. The rank of each cell is also displayed by a black bar plot and the vertical dashed black line represents the average rank. The dendrogram on the left represents strain ordered by hierarchical clustering.

Finally, we investigated the strain phenotypic variability according to the environmental conditions. This characteristic is very important in enology since industrial strains might be used in different grape musts with contrasted physicochemical properties. Therefore, the assessment of phenotypic robustness of industrial starters is crucial for optimizing their use in a wine making process. We computed the phenotypic variance of the 35 strains by using the non-normalized dataset. The overall results are shown on Fig 8, panel A. Strains showing a low variance value (green tones) had similar phenotypic behavior in the 5 grape musts. On the contrary, high variance (red tones) values indicated a contrasted phenotypic response according to the must. Some industrial strains such as C23, C10 or C12 showed a strong robustness to environmental change. In contrast, the monosporic clones SB, GN and M2, as well as some commercial strains (C22, C7, C18) appeared to be quite sensitive to environmental changes (high phenotypic variability indicated by red tones). The source of the lack of robustness was investigated by splitting the 35 strains in two groups according to their phenotypic robustness (variance). The less robust quartile was compared to the 75 % more robust strains in the 5 grape juices. Thus the conditions that generate a lack of robustness could be identified. For example, *lp* was only significantly different for the two groups only in SB15 (3.2 time longer for the non-robust group) (Fig 8, panel B). In this example, identified grape must had the strongest initial SO_2_ concentration (67 mg.L^-1^), which is known to strongly affect the lag phase [8]. All the strains of the non-robust group (C1, C11, F15, C15, M2, C22, SB, C31) are therefore not suitable for running fermentations in highly sulphited grape musts. This is also the case for another group of strains (C4, C7, C17, GN, C18, C21, C24, C25, C27), which only produced high concentrations of *SO*_*2*_ at the end of the fermentation in SB15 (Fig 8, panel B). The two Merlot grape musts (M14 and M15), which are harsh to ferment, were those that best discriminated the strains for the *t80* and *pyruvate* robustness (S4 Fig). For *acetic acid,* SB14 mainly increased the variance of the less robust strains. For example, the strains C18 and C24 produced high levels of *acetic acid* in white grape musts but they showed a moderate production in the three red grape musts. This result suggests that these two strains are not suitable for white grape musts. Finally, due to its higher sugar concentration, CS15 promoted high glycerol production and exacerbated differences between strains (S4 Fig).

**Fig 8.**
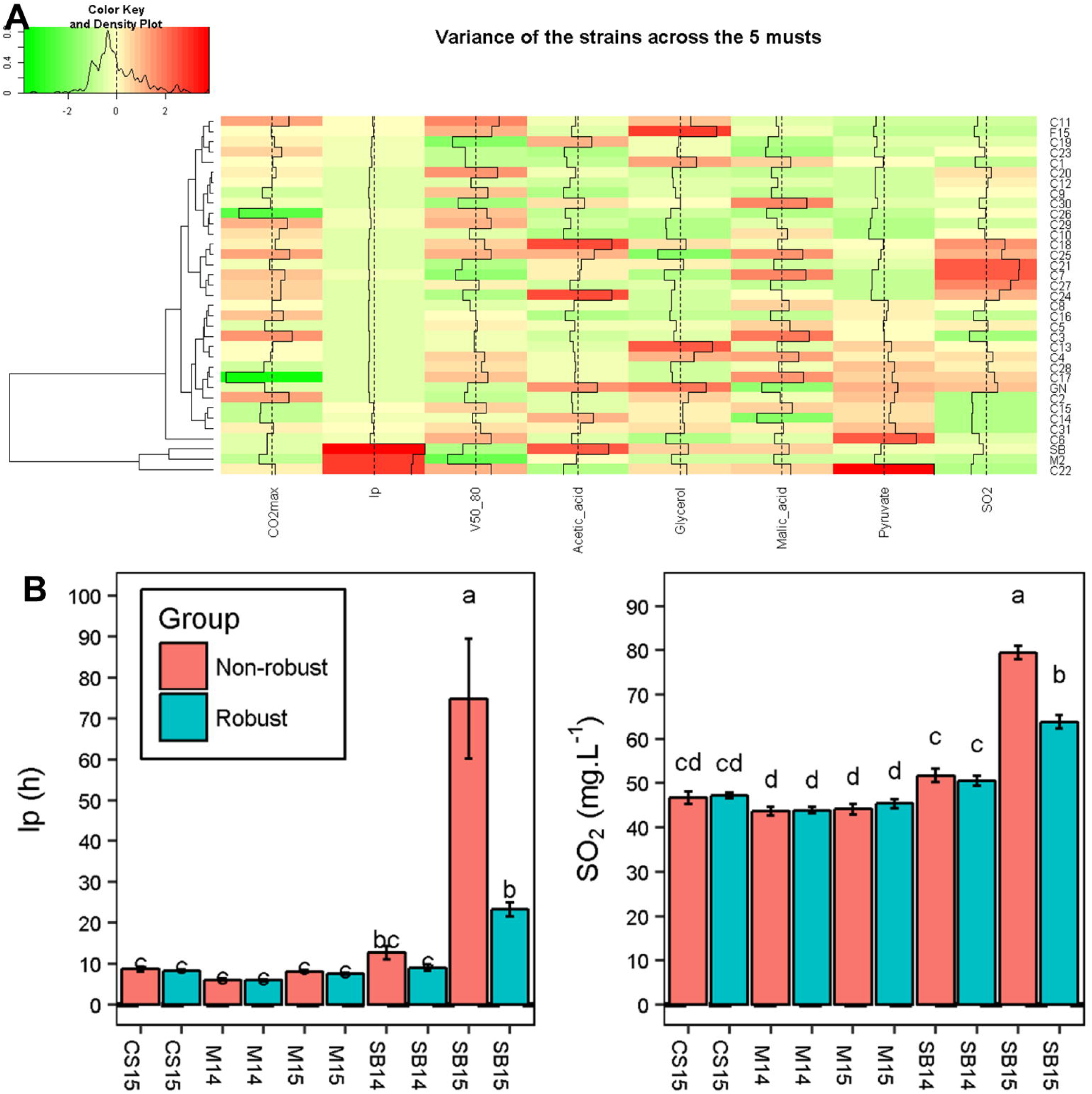
Phenotypic variance of 35 strains in 5 grape juices. Panel A. For each strain, the variance was computed for the 5 average phenotypic values in the 5 grape musts. Variance is scaled by column and its level is represented by a color palette from green (lowest variance) to red (highest variance) as displayed by the color key. The value of each cell is also displayed by black bar plots and the vertical dashed black lines represents the average variance. Strains are ordered by hierarchical clustering that is represented by the dendrogram on the left. Panel B. Comparison of *lp* and *SO*_*2*_ between robust and non-robust strains according to grape musts. The data shown are the mean of 8 strains (non-robust group) or 27 strains (robust group), the error bars represent the standard error. Different letters indicate significant differences between groups (Tukey's honest significant difference test, significance level, α = 0.05).

## Discussion

### A new platform for measuring quantitative traits related to wine fermentation

The wide development of NGS technologies gives the opportunity to collect large sets of genomic data that could be used for dissecting the genetic architecture of complex traits using both QTL mapping and GWAS approaches [15]. In order to implement genetic studies efficiently, this genomic data must be completed with massive sets of phenotypic data. The high throughput measurement of phenotypes is therefore a crucial point for finding out new genetic determinisms. In the last decade, the term of “phenomics” has been used to describe methods aiming at measuring phenotype at a large-scale [24]. Mostly based on the measurement of OD [45] or plate growth [46], the parallel measurement of basic growth parameters in numerous media can be performed. Although this approach is very useful for screening growth-related phenotypes, other complex traits of industrial interest, such as fermentation kinetics and end-product metabolites can neither be measured in micro-plates nor in agar plates.

In this study, we set up a standardized method for assessing alcoholic fermentation experiments at a relatively large scale (>300 samples per batch). By reducing the fermentation volume to 5 mL in standard SV, we conserved a very accurate estimation of fermentation kinetics that matches well with the methods previously used [47]. Here, the fermentation time course was followed manually by weighing each SV two times per day with a precision balance. However, robotic solutions for an automatic handling of the SV could easily be implemented thanks to the standardized format of the vials used. In order to face the large sample analysis set required, we successfully coupled our fermentation setup with a robotized enzymatic platform for measuring eight enological metabolites in 1 mL samples. Unfortunately, we failed to efficiently measure ethanol, since the enzymatic kit used was not sufficiently accurate for high ethanol concentrations. Alternatively, the estimation of total CO_2_ loss was very precise (average CV<3 %) and perfectly matched with the production of ethanol during the alcoholic fermentation [34]. During this study, we also demonstrated that at the end of the alcoholic fermentation, many volatile compounds produced by yeast metabolism could be readily analyzed by GC-MS after an automated solid-phase micro-extraction [31]. Coupling analytical facilities and developing robotic handling of SV will be the next steps for developing large screening programs.

### Assessment of some GxE interactions relevant in enology

Although the volume fermented is far from representing those of vats used during industrial wine production, our setup was close as possible to the enological conditions. The effects of some parameters that are relevant for enology (grape must, strain, micro-oxygenation level) could be tested. First of all, we used natural grape musts rather than synthetic media that might be less pertinent for assessing quantitative traits due to their incomplete composition [20,48]. As previously demonstrated, frozen grape juices conserved their fermentation properties and can be kept for long periods [29]. Moreover, in this work we only tested a panel of commercial starters that are used in various geographic areas for the production of red, white, *rosé* and sparkling wines. This contrasts with previous studies that also included *S. cerevisiae* strains from other origins [49,50]. By using only commercial starters, we captured here a phenotypic variability having an industrial relevance and reflecting those proposed to the winemaker. Finally, the shaking of SV was able to mimic micro-oxygenation in a reproducible manner. The amount of oxygen transferred during the 20 first hours (2-4 mg/L^-1^ of O_2_) is close to that occured in red winemaking practices [37]. Although the micro-oxygenation is provided by several pumping-over operations in the cellar, we were able to reproduce this effect in our small design vessels with similar scale values. This was confirmed by observing effects that are similar to those already known in enological practices. Indeed, a higher level of micro-oxygenation accelerates the fermentation rate [37,51,52], decreases the production of acetic acid [41,51], and increases the production of glycerol [38,39,51,53]. The shaking conditions also had an impact on the stripping of volatile molecules such as SO_2_.

Interestingly, by assaying 32 volatile compounds using a GC-MS approach we demonstrated that shaken conditions do not impact all the volatile molecules in the same way suggesting that the oxygen transfer could influences the production of aromatic compounds and in particular esters. Unravelling the impact of oxygen on esters production during the alcoholic fermentation is not trivial. According to the quantity and the addition moment, the oxygen effect may indeed be drastically different. The oxygen supplementation of grape must in winemaking conditions resulted in an increase of the concentration of higher alcohol acetates and branched chain ethyl esters, and in a decrease of fatty acid ethyl esters [51,54]. Aside higher alcohol acetates that were 2 times higher in non-shaken conditions, our findings are broadly in agreement with previous data measured in a cellar [51,54]. The similar response between 5 mL SV and vats of several liters is very encouraging and demonstrates that our setup could be relevant for assessing the aromatic production of a large set of strains/conditions. The change in proportion of 2-phenylethyl acetate to ethyl-phenylacetate could be a signature of micro-oxygenation, as the proportion of the most oxidized ester (ethyl-phenylacetate) is greater with agitation. Moreover, the relative higher production of acetate of higher alcohols in non-shaken conditions could be explained by the fact that the moment of oxygen addition and metabolizing is drastically different between the yeast growth and stationary phase [35,55]. For example, in a brewing context, when oxygen is added during the fermentation, a decreased production of higher alcohol acetates can be observed [55,56], thus supporting our observations (S5 Table). Conversely, oxygen addition has been reported to increase the concentration of ethyl esters and to reduce the concentration of acetate esters and higher alcohols [57]. These seemingly contradictory results can be also due to strain-by-oxygenation interactions. Indeed 16 of the 32 compounds assayed showed *strain x environment* interactions. This setup thus could be useful in the future to better investigate the physiological and enological consequences of micro-oxygenation for up to very large panels of yeast strains.

Thanks to this setup, we gained insight on other GxE interactions between wine strains and environmental conditions. For example, GN maintained a constant level of *acetic acid* in M15, regardless of the level of micro-oxygenation. This particular feature, which is a relevant trait in enology, suggests that acetic acid metabolism is poorly impacted by hypoxia in this strain. Another interaction was observed for the strain M2, for which the long lag phase observed in sulphited grape must (SB14) is reduced by the micro-oxygenation. Those preliminary observations open perspectives for studying the phenotypic response of yeast strains to micro-oxygenation at a large scale.

### Survey of the fermentation performances of 35 enological strains in 5 grape musts

As a matter of proof, we measured the phenotypic performances of 35 strains including 31 industrial starters. After 3 weeks of fermentation, we measured in the same batch 11 traits in 5 different grape juices (350 fermentations), supporting the efficiency of the method for high throughput phenotyping. We have observed an important grape must effect on the phenotypes (Table 3). This effect was generated by the basic physicochemical characteristics of the grape musts (concentration of sugar, malic acid and SO_2_ *etc*.). In order to go beyond this effect, the response of each strain was normalized according to the grape juice. Therefore the principal effect of the media was eliminated allowing the comparison of each strain’s response measured in five grape juices. With the normalized dataset, we first investigated the relations between the quantitative traits measured for 35 strains. The large panel of commercial strains and the measurement of each trait in 5 conditions reinforces the robustness of these links and ensures the generalization of the conclusions that can be drawn. Omitting the obvious correlations found between kinetic parameters, we shed light on other correlations that can reflect important metabolic trade-offs. A strong correlation was identified between *malic acid* and kinetic parameters (*T35*, *T50* and *T80*). Thus, we found that fast fermenting strains were also those consuming more malic acid. This link had already been observed for the ML01 strain, which has been genetically modified to carry out the malolactic fermentation [42]. ML01 has a higher fermentation rate than the parental strain. This can be explained by the deacidification of the media caused by the malic acid consumption that can provide more permissive pH conditions. However, it is important to note that this effect appears to occur in low pH conditions. Therefore, as the pH of the grape musts used in our study ranged from 3.19 to 3.58, other mechanisms were probably involved. For example it is known that malic acid plays an important role in carbon metabolism. During fermentation its decarboxylation provides pyruvate which could play an anaplerotic effect on biomass and/or on ethanol synthesis [58,59]. A second positive correlation was found between the duration of the lag phase (*lp*) and the glycerol production, suggesting that strains starting the fermentation later produce more glycerol. The production of glycerol at the beginning of fermentation helps resoring the redox balance by regenerating the NAD^+^ consumed via the reaction catalyzed by glyceraldehyde-3-phosphate dehydrogenase at the beginning of the glycolysis [60]. Indeed, at that stage, the regeneration of NAD^+^ by alcohol dehydrogenase is subject to inhibition by the formation of complexes between acetaldehyde and bisulfites ions. Thus, strains that are able to rapidly start the alcoholic fermentation do not need to produce high glycerol amounts to compensate for this NAD + deficiency. Moreover, glycerol production is a well-known response to osmotic stress, which results from the high sugar concentration found in grape juice [61]. As osmotic shock affects cell growth and the lag phase [62], the high producer strains could be more adapted to initiate the alcoholic fermentation promptly after inoculation.

This dataset was also used to evaluate and compare the performance of the strains. This comparison revealed groups of strains with distinct phenotypic profiles. This disparity shows that despite the high specialization level of wine starters [63], the completion of the fermentation takes place over a wide range of production or consumption of important end-products. For example, groups C and D defined in Fig 7 mainly discriminate the strains having the highest fermentation rate by their *glycerol* and *SO*_*2*_ production levels. This suggests that high performing (commercial) strains adapted to winemaking conditions have undergone different adaptive strategies that have modelled their central metabolism in order to accomplish the alcoholic fermentation. Strain robustness against the grape must parameter was evaluated, leading to the identification of the most robust commercial starters. Robustness is a critical factor for the wine industry, as it ensures successful fermentations in a wide range of grape musts. Some grape musts with extreme characteristics (SO_2_ or sugar concentrations) highlighted the weakness of the less robust ones leading to the identification of the type of grape must for which they are the most suited. The setup developed in the present study could help to identify the physicochemical factors (amino acids, vitamins, cofactors or polyphenols) that could be a source of inappropriate phenotypic responses. The identification of enological factors that affect the performance of strains is of great interest. It has already been shown for example that the effect of temperature during fermentation was dependent on the strain used [25]. The fermentation system implemented here is well adapted to push forward the identification of new factors of this type.

## Acknowledgments

The authors want to thank Pr Vladimir Jiranek for kindly giving some strains used in this work. We also thank Warren Albertin, Brendan Smith, Marina Bely and Joana Coulon for their correction that helped improving the manuscript. And finally, we thank Elodie Kaminski that helped managing fermentations.

## Supporting information

**S1 Fig SV setup**

On the left, a SV filled with 5 mL of grape juice (SB14) and with a hypodermic needle to allow the CO_2_ release. On the right 70 vials on a rack illustrating the possibility of managing hundreds of fermentations in parallel.

**S2 Fig Oxygen impact on ester production**

Panel A. The data shown are the mean proportion of PhC2C2 to C2PhC2 of the 4 strains in 2 replicates, the error bars represent the standard error. Different letters indicate significant differences between groups (Tukey's honest significant difference test, significance level, α = 0.05).Panel B. The data shown are mean of 2 replicates, the error bars represent the standard error. Different letters indicate significant differences between groups (Tukey's honest significant difference test, significance level, α = 0.05). Table represents ANOVA results (pval, and % of variance explained).

**S3 Fig Correlations between traits**

Scatter plots of correlated traits. Each dot represent the average phenotypic values of a strain across the 5 grape must from the normalized dataset. The blue line represents the linear regression line and the shaded area represents the confidence interval of the regression (0.95).

**S4 Fig Comparison of the phenotypic values between robust and non-robust strains according to grape musts.**

The data shown are the mean of 8 strains (non-robust group) or 27 strains (robust group), the error bars represent the standard error. Different letters indicate significant differences between groups (Tukey's honest significant difference test, significance level, α = 0.05).

**S1 Table yeast strains used**

**S2 Table Dilution and volume of sample used for robotic enzymatic assay**

**S3 Table List of the 32 esters analyzed**

**S4 Table SB14 dataset**

Data presented are the mean of six fermentation replicates of SB 14 grape must. The residual sugars (glucose + fructose) at the end of the fermentation was not shown and was always lower than 1.5 g.L^-1^. Statistical differences within strains and modalities was assayed by Tukey's honest significant difference test, significance level, α = 0.05, the different groups were shown by a letter code: groups sharing the same letter are non-significantly different.

**S5 Table Esters dataset**

**S6 Table Micro-oxygenation and grape must interaction dataset**

**S7 Table phenotypic data of 35 commercial strains in 5 grape juices (raw data)**

**S8 Table phenotypic data of 35 commercial strains in 5 grape juices (centered reduced data)**

